# Ivermectin increases striatal cholinergic activity to facilitate dopamine terminal function

**DOI:** 10.1101/2023.11.30.569432

**Authors:** Hillary A. Wadsworth, Alicia M.P. Warnecke, Joshua C. Barlow, J. Kayden Robinson, Emma Steimle, Joakim Ronström, Pacen Williams, Christopher Galbraith, Jared Baldridge, Michael W. Jakowec, Daryl L. Davies, Jordan T. Yorgason

**Author notes:** **Corresponding Author:** Jordan T. Yorgason, PhD.

## Abstract

Ivermectin (IVM) is a commonly prescribed antiparasitic treatment with pharmacological effects on invertebrate glutamate ion channels resulting in paralysis and death of invertebrates. However, it can also act as a modulator of some vertebrate ion channels and has shown promise in facilitating L-DOPA treatment in preclinical models of Parkinson’s disease. The pharmacological effects of IVM on dopamine terminal function were tested, focusing on the role of two of IVM’s potential targets: purinergic P2X4 and nicotinic acetylcholine receptors. Ivermectin enhanced electrochemical detection of dorsal striatum dopamine release. Although striatal P2X4 receptors were observed, IVM effects on dopamine release were not blocked by P2X4 receptor inactivation. In contrast, IVM attenuated nicotine effects on dopamine release, and antagonizing nicotinic receptors prevented IVM effects on dopamine release. IVM also enhanced striatal cholinergic interneuron firing. L-DOPA enhances dopamine release by increasing vesicular content. L-DOPA and IVM co-application further enhanced release but resulted in a reduction in the ratio between high and low frequency stimulations, suggesting that IVM is enhancing release largely through changes in terminal excitability and not vesicular content. Thus, IVM is increasing striatal dopamine release through enhanced cholinergic activity on dopamine terminals.

## Introduction

Mesostriatal circuitry includes major inputs from dopamine (DA) neurons originating in the substantia nigra compacta (SNc) that synapse onto local medium spiny neurons to modulate output from the direct and indirect striatal pathways. Local regulators of this circuit include large aspiny cholinergic interneurons (CINs) that fire rhythmically due to intrinsic activity(Bennett & Wilson, 1999), and are further activated by glutamate (e.g. thalamic inputs)(Mamaligas, Barcomb, & Ford, 2019), and by hyperpolarization activated currents driven by GABA inputs(Brown et al., 2012). Cholinergic firing can also depolarize DA terminals through activation of nicotinic acetylcholine receptors (nAChRs), resulting in cholinergic-evoked DA release or modulation of ongoing release.(Brundage et al., 2022; Rice & Cragg, 2004; Wadsworth et al., 2023; Yorgason et al., 2022; Yorgason, Zeppenfeld, & Williams, 2017; Zhang & Sulzer, 2004; Zhou, Liang, & Dani, 2001) Alterations in DA inputs and related cholinergic activity are known to drive changes in movement and mood, and underlie related disorders such as Parkinson ’s disease.(Aarsland, Påhlhagen, Ballard, Ehrt, & Svenningsson, 2011; Chinta & Andersen, 2005; Miller & O’Callaghan, 2015)

The present study examines the interactions between striatal DA and cholinergic local circuits and the commonly prescribed anti-parasitic ivermectin (IVM). IVM is known to affect many DA associated behaviors, including ethanol consumption, anxiety, sensorimotor deficits, and sociocommunicative behavior(Franklin et al., 2014; Khoja, Asatryan, Jakowec, & Davies, 2019; Khoja et al., 2018; Khoja et al., 2016; Wyatt et al., 2013; Yardley et al., 2012). Indeed, IVM has also been shown to improve L-DOPA induced behaviors, including in preclinical Parkinson’s disease (PD) animal models.(Khoja et al., 2016; Warnecke, Kang, Jakowec, & Davies, 2020) However, no studies have been conducted to determine whether IVM influences striatal DA release. Therefore, fast scan cyclic voltammetry (FSCV) was used to measure synaptic DA release in the dorsal striatum (DS) and pharmacological agents were applied during experiments to elucidate the mechanistic effects of IVM on DA release.

## Methods

### Animal Subjects

Female and male C57BL/6 (>30-d-old) were bred and cared for in accordance with the National Institutes of Health Guide for the Care and Use of Laboratory Animals. Animals were housed on a reverse 12:12 h light/dark cycle (lights on from 10 PM to 10 AM) in groups of 2–5/cage and given ad libitum access to food and water. For imaging experiments, Macrophage Fas-Induced Apoptosis (MaFIA) transgenic mice (RRID:IMSR_JAX:005070) were used to image microglia. Experimental protocols were approved by the Brigham Young University Institutional Animal Care and Use Committee according to the National Institutes of Health *Guide for the Care and Use of Laboratory Animals*.

### Brain Slice Preparation

Coronal brain slices were obtained as previously described(Brundage et al., 2022; Yorgason et al., 2017). Briefly, animals were anesthetized with isoflurane (5%), decapitated, and brains were rapidly dissected and sectioned into 220 μm slices in artificial cerebrospinal fluid (ACSF) cutting solution. The ACSF cutting solution (pH = ∼7.4) was oxygenated at 95% O_2_/5% CO_2_ and consisted of (in mM) 126 NaCl, 2.5 KCl, 1.2 NaH_2_PO_4_, 2.4 CaCl_2_, 1.2 MgCl_2_, 21.4 NaHCO3, 11 glucose and 0.1 ketamine (for glutamate receptor blockade). Slices were transferred to a recording chamber with continuous ACSF flow (2.0 mL/min) maintained at 34–36 °C. The dorsal striatum was visualized at the level of the dorsal horn under low magnification with Nikon Diaphot inverted microscopes in the transmitted light mode and Olympus X51 microscopes with transmitted infrared Dodt gradient contrast imaging.

### Fast scan cyclic voltammetry recordings

Electrically evoked DS DA release was obtained using FSCV. Carbon fiber electrodes (CFEs) were Nafion coated using a 1.5V 90s electrodeposition pretreatment for L-DOPA experiments(Qi et al., 2016). Dopamine release was electrically evoked every 2 min by monophasic stimulation from a KCl-filled micropipette placed 100–200 μm from the CFE. L-DOPA experiments used an alternating single pulse/5 pulse stimulation protocol (0.5 msec pulse, 350 μA, 20Hz). Additional experiments used a frequency stimulation protocol that included 5Hz, 20Hz, and 100Hz 5 pulse stimulations. Experiments performed with hexamethonium used a 4 msec pulse to account for the signal disruption with a shorter 0.5 msec pulse. The CFE potential was linearly scanned from −0.4 to 1.2 V and back to −0.4 V vs Ag/AgCl (scan rate = 400 V/s). Cyclic voltammograms were recorded every 100 msec (10Hz) with ChemClamp potentiometers (Dagan Corporation, Minneapolis, MN, USA) or inhouse developed potentiostats. Recordings were performed and analyzed using Demon Voltammetry software as described below(Yorgason, España, & Jones, 2011).

### Drug preparation and administration

IVM (cat. no. NDC 55529-012-01, Norbrook Laboratories, Ltd, Newry, North Ireland, UK), Nicotine (cas. no. 54-11-5, Sigma-Aldrich, St. Louis, Missouri, USA), Hexamethonium bromide (cas. no. 55-97-0, Cayman Chemical Company, Ann Arbor, Michigan, USA), L-DOPA (cat. no. PHR1271, Sigma-Aldrich, St. Louis, Missouri, USA), and 5-(3-Bromophenyl)-1,3-dihydro-2H-benzofuro[3,2-e]-1,4-diazepin-2-one (5-BDBD) (cat. no. T22518, TargetMol, Wellesley Hills, Massachusetts, USA) were dissolved in stock solutions and then diluted into ACSF at specified concentrations (50 µM IVM, 300 nM Nicotine, 200 µM Hexamethonium, 10 µM L-DOPA, and 10 µM 5-BDBD). Drugs brain slice administration used either gravity-based flow system or peristaltic pumps (1-2 ml/min).

### Multiphoton Imaging

A transgenic mouse expressing enhanced green fluorescent protein (EGFP) on the colony stimulating factor 1 receptor on a C57BL/6J background (Macrophage Fas-Induced Apoptosis, MAFIA) was used to visualize microglia. The animal was anesthetized in 2-4% isoflurane and sacrificed as described above. The brain was extracted, and slices were prepared as described above, targeting the dorsal striatum. The slice was then immediately put into ACSF to incubate with the P2X4 antibody (50:1; Thermo Fisher Scientific, PA5-37880; AF647 conjugated) at room temperature in the dark for 30 minutes. The brain slice was moved to the two-photon recording chamber and immersed in ACSF (flow rate of ∼1-2 ml/min), to wash away excess antibody. The custom in-house built two-photon microscope used a Ti-Sapphire Chameleon Discovery NX Laser (Coherent) tuned to 870 nm (optimized for simultaneous excitement of EGFP and Alexa Fluor 647). Fluorescent emission was visualized using a 40x (0.8NA) water immersion objective (Olympus). A Z-stack was collected with 3 μm between slices, with a total range of 17 slices for simultaneous detection of GFP (520 nm cleanup filter) and Alexa Fluor 647 (670 nm cleanup filter). Images were combined and colorized using Fiji(Schindelin et al., 2012).

### Cell-attached electrophysiology recordings in brain slices

Electrophysiology studies utilized borosilicate glass capillary electrodes (2.5–6 MΩ). For cell-attached recordings of CIN firing (NaCl 150 mM inside the pipette), a seal (10MΩ – 1GΩ) was created between the cell membrane and the recording pipette. Spontaneous spike activity was then recorded in voltage-clamp mode with an Axon Instruments Multiclamp 700B (Molecular Devices, San Jose, CA, USA) amplifier and sampled at 3 kHz using an Axon 1440A digitizer (Molecular Devices) and collected and analyzed using Mini Analysis (Synaptosoft: Decatur, GA) and/or Axograph 10 (Axograph, Sydney, Australia). A stable baseline recording of current activity was obtained for 5 min before perfusion of Ivermectin (50 μM) which was applied for ∼10 minutes.

### Statistical Analysis

Dopamine release experiments were analyzed in Demon Voltammetry(Yorgason et al., 2011). To compare across multiple animals and slices, DA release signals were averaged (within subject) across the last three 1 pulse or 5 pulse ACSF recordings to establish a baseline value that subsequent DA signals were normalized to. Thus, post-drug conditions were compared to pre-drug baseline conditions in the same slice for a within-subject experimental design. Electrophysiology CIN firing rate experiments used Axograph and Minianalysis for analysis of action potentials and experiments were also performed using a within-subject design. Statistical analysis was performed using Prism 5 (GraphPad). Significance for all tests was set at *p* < 0.05. Values are expressed as mean ± SEM. Significance levels are indicated on graphs with asterisks *,**,***, corresponding to significance levels p < 0.05, 0.01 and 0.001, respectively. Data will be shared on a per request basis.

## Results

### IVM enhances DA release independent of P2X4 receptor activation

Bath application of IVM (50 µM) results in increased electrically evoked DS DA release, as measured by FSCV (**Fig. 1A**-**B**). IVM is a known positive allosteric modulator (PAM) of P2X4 receptors(Khoja et al., 2016; Priel & Silberberg, 2004). The P2X4 receptor is expressed in the DS, in particular on microglia, but also on non-microglia cells (**Fig. 1C**). Co-application of the P2X4 receptor antagonist 5-BDBD (10 µM) did not prevent IVM induced DA increases (**Fig. 1D**; one-way ANOVA; *F*_(2,41)_ = 10.32, p = 0.0002). Dopamine release can be sensitive to local circuit activity that manifests as increases in release across stimulation frequencies.(Schilaty et al., 2014; Yorgason, Rose, McIntosh, Ferris, & Jones, 2015; Yorgason et al., 2022) Therefore, a high to low stimulation frequency protocol was used to examine IVM and P2X4 receptor interactions on DA release circuits. Generally, neither IVM nor 5-BDBD affected the frequency response curve compared to baseline (**Fig. 1E**; two-way ANOVA; drug, *F*_(2,164)_ = 2.166, p = 0.1179; frequency, *F*_(3,164)_ = 0.7402, p = 0.5295; interaction, *F*_(6,164)_ = 0.3469, p = 0.9109). The frequency response area under the curve (AUC) remained unchanged between the different treatments (**Fig. 1F**; one-way ANOVA; *F*_(2,41)_ = 0.5950, p = 0.5563). Together, this data indicates that IVM-induced increases in DS DA release are independent of P2X4 receptor activation and that IVM has no apparent effect on the DA frequency response.

**Figure 1:**
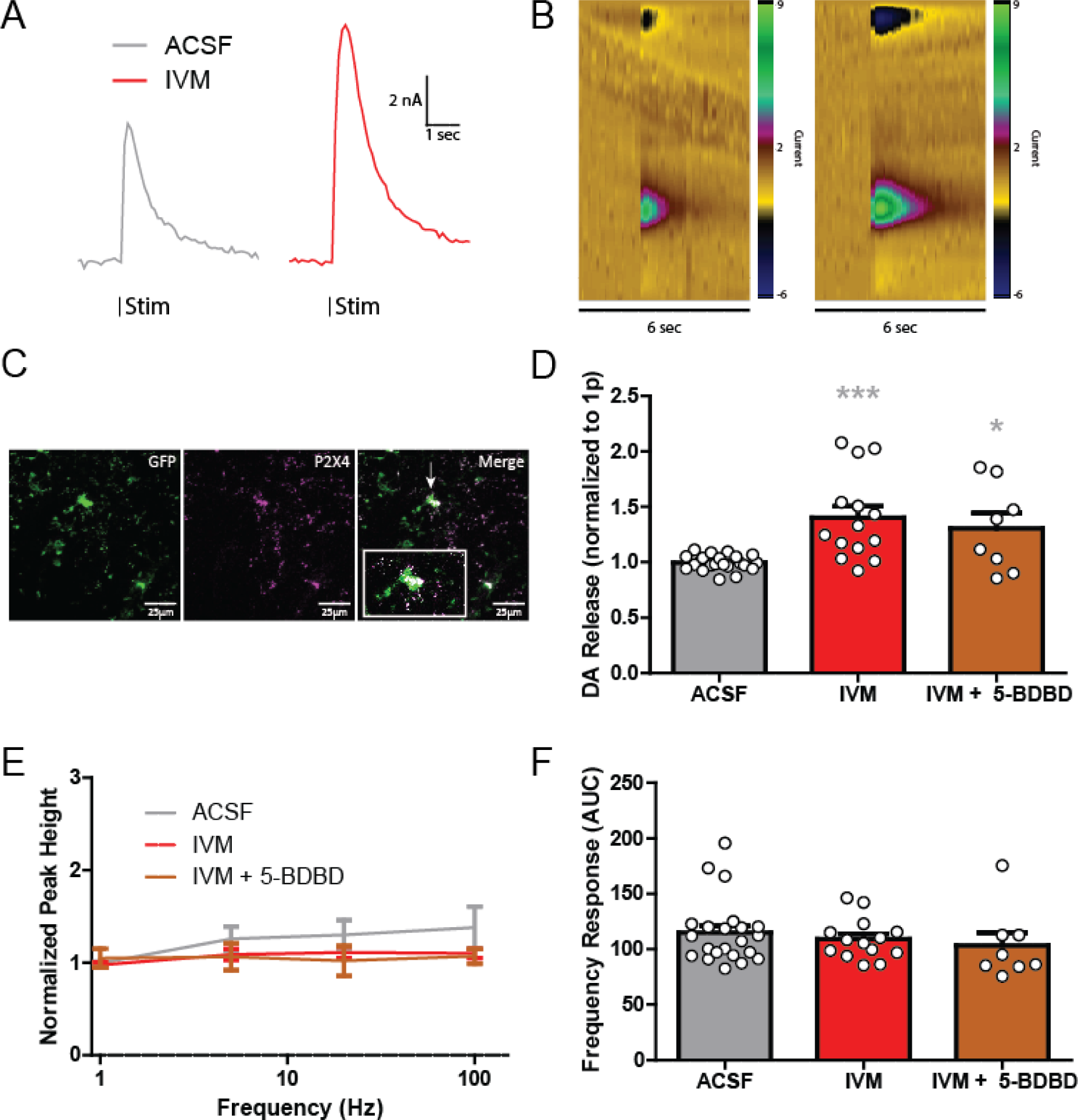
IVM enhances Dopamine Release independent of P2X4 receptor activation. **A** Representative traces of single pulse electrical evoked dopamine release in the DS before (left) and after (right) IVM application. **B** Representative color plots of dopamine release before (left) and after (right) IVM application. **C** Image showing P2X4 receptor co-localization on microglia in the DS. An antibody against P2X4 receptors was used along with Macrophage Fas-Induced Apoptosis (MaFIA) transgenic mice with GFP labeled microglia. **D** Ivermectin, as well as IVM with 5-BDBD increases single pulse dopamine release in the dorsal striatum compared to normalized ACSF release. **E** Ivermectin, as well as IVM with 5-BDBD does not affect DA release at multiple frequencies compared to ACSF alone. **F** The overall frequency response, as measured by area under the curve (AUC) of the previous figure (D), is not affected by IVM or IVM with 5-BDBD. Asterisks *,*** indicate significance levels p < 0.05 and p < 0.001 respectively compared to ACSF pre-treatment.

### IVM attenuates nicotine effects on DA release

IVM is a known nAChR PAM,(Collins & Millar, 2010; Krause et al., 1998) and nAChRs are powerful activators and modulators of DA terminal activity.(Yorgason et al., 2017) Further, nAChRs PAMs, such as alcohol, increase DA terminal function resulting in greater release.(Gao et al., 2019) Accordingly, experiments were conducted to assess if IVM modifies known nAChR effects on DA release. Cholinergic interneurons release acetylcholine to directly depolarize DA terminals(Wadsworth et al., 2023; Zhou et al., 2001). However, prolonged activation of nicotinic receptors results in alterations in electrically evoked DA release which are often described as a high-pass filter, and release becomes biased towards high frequency stimulation conditions(Rice & Cragg, 2004) (**Fig. 2A**). IVM itself did not strongly exhibit high pass filter properties (**Fig. 2B**). Next, experiments were conducted to test if IVM influenced nicotine’s effects on DA release (**Fig. 2**). In contrast to IVM facilitatory effects on single pulse stimulated DA release, nicotine alone reduced DA release. However, concurrent application of IVM (50 µM) and nicotine (300 nM) resulted in increased variability and significantly impaired nicotine inhibitory effects on DA release (**Fig. 2C**; one-way ANOVA; *F*_(3,45)_ = 9.944, p < 0.0001). Furthermore, nicotine alone enhanced the 20Hz 5:1 pulse ratio (**Fig. 2D**). However, IVM and nicotine co-application resulted in a reduction in this ratio similar to baseline and IVM alone (**Fig. 2D**; one-way ANOVA; *F*_(3,45)_ = 11.11, p < 0.0001). Examining the full frequency stimulation condition curve, the well characterized high-pass filter effect was observed with nicotine (**Fig. 2A**,**E**). However, this effect was greatly diminished by IVM co-application, and IVM alone had no apparent effect on the frequency response (**Fig. 2A**-**B**,**E**; two-way ANOVA; drug, *F*_(3,209)_ = 30.97, p < 0.0001; frequency, *F*_(3,209)_ = 25.48, p < 0.0001; interaction, *F*_(9,209)_ = 10.37, p < 0.0001). IVM appears to block or counter the effects of nicotine on DA release from single pulse and multiple pulse high frequency stimulations. This finding is summarized in the AUC measures from the stimulation frequency response curve (**Fig. 2F**). The inclusion of IVM with nicotine significantly reduced the AUC magnitude compared to nicotine alone (**Fig. 2F**; one-way ANOVA; *F*_(3,45)_ = 19.93, p < 0.0001). Thus, IVM enhances DA release on its own and opposes the nicotine-induced high-pass frequency bias, suggesting that IVM effects on DA release may involve effects on the acetylcholine system.

**Figure 2:**
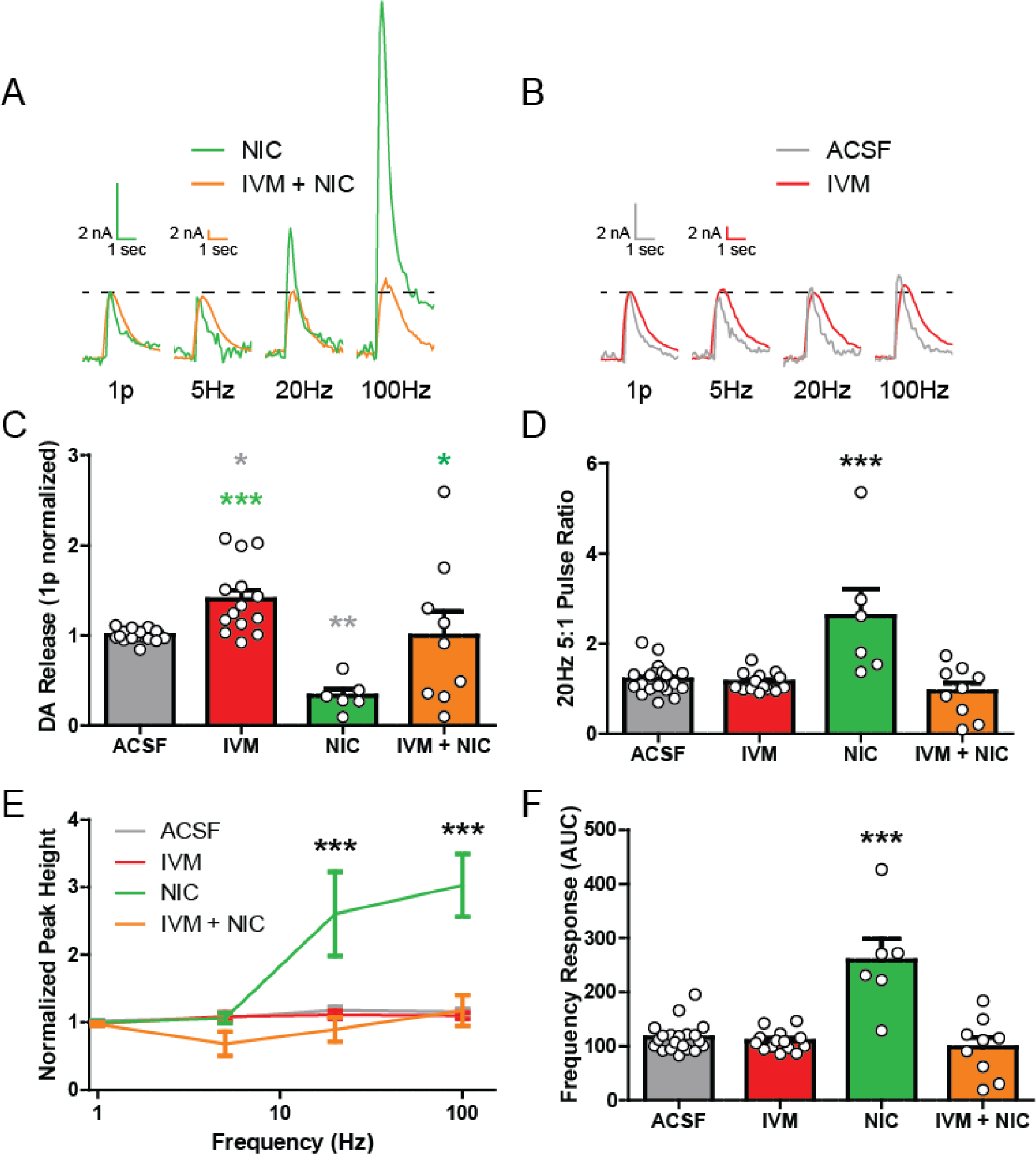
IVM attenuates Nicotine effects on DA release. **A** Representative traces of electrically evoked dopamine release in the DS at multiple frequencies. Comparing Nicotine (NIC) and IVM + NIC, each trace is normalized to 1 pulse release (dotted line). **B** Representative traces of DS dopamine release at multiple frequencies comparing ACSF and IVM. Each trace is normalized to 1 pulse release (dotted line). **C** Ivermectin increases single pulse dopamine release while NIC decreases dopamine release. Ivermectin attenuates the effect of NIC on single pulse dopamine release. **D** Nicotine alone increases the 20Hz 5 pulse ratio compared to ACSF, IVM, and IVM + NIC. **E** Nicotine alone shows increased dopamine release in response to high frequency, 5 pulse stimulations, with AUC represented in **F**. Asterisks *,** indicate significance levels p < 0.05 and p < 0.01 respectively compared to ACSF pre-treatment. Asterisks *,*** indicate significance levels p < 0.05 and p < 0.001 respectively compared to nicotine treatment. Asterisks *** indicate significance levels p < 0.001 compared to all other treatments.

### IVM attenuation of nicotine effects is not through P2X4 receptors

IVM-mediated DA enhancement is not through IVM-PAM effects on P2X4 receptors. However, nicotine may be recruiting other circuit effects that could reveal a role for P2X4 receptors. Thus, additional experiments were performed to test if P2X4 receptors are involved in IVM reductions in nicotine frequency response effects on DA release (**Fig. 3**). If IVM is modulating the nicotinic effect through P2X4 receptor interactions, then presence of the P2X4 antagonist 5-BDBD should restore nicotinic effects on DA release. Experiments with IVM, nicotine, and 5-BDBD applied simultaneously revealed a similar effect to ACSF alone and was significantly different from nicotine alone (**Fig. 3A**,**C**; one-way ANOVA; *F*_(3,42)_ = 6.779, p = 0.0008). Similarly, nicotine effects on the 5:1 pulse ratio were not restored with 5-BDBD application (**Fig. 3B**,**D**; one-way ANOVA; *F*_(3,42)_ = 10.64, p < 0.0001). 5-BDBD did not restore the nicotine high frequency DA release effects (**Fig. 3E**; two-way ANOVA; drug, *F*_(3,163)_ = 24.71, p < 0.0001; frequency, *F*_(3,163)_ = 18.82, p < 0.0001; interaction, *F*_(9,163)_ = 8.085, p < 0.0001), which is summarized with the AUC measures from the frequency response (**Fig. 3F**; one-way ANOVA; *F*_(3,40)_ = 17.98, p < 0.0001). Therefore, IVM effects on the cholinergic system are independent of P2X4 receptor activation.

**Figure 3:**
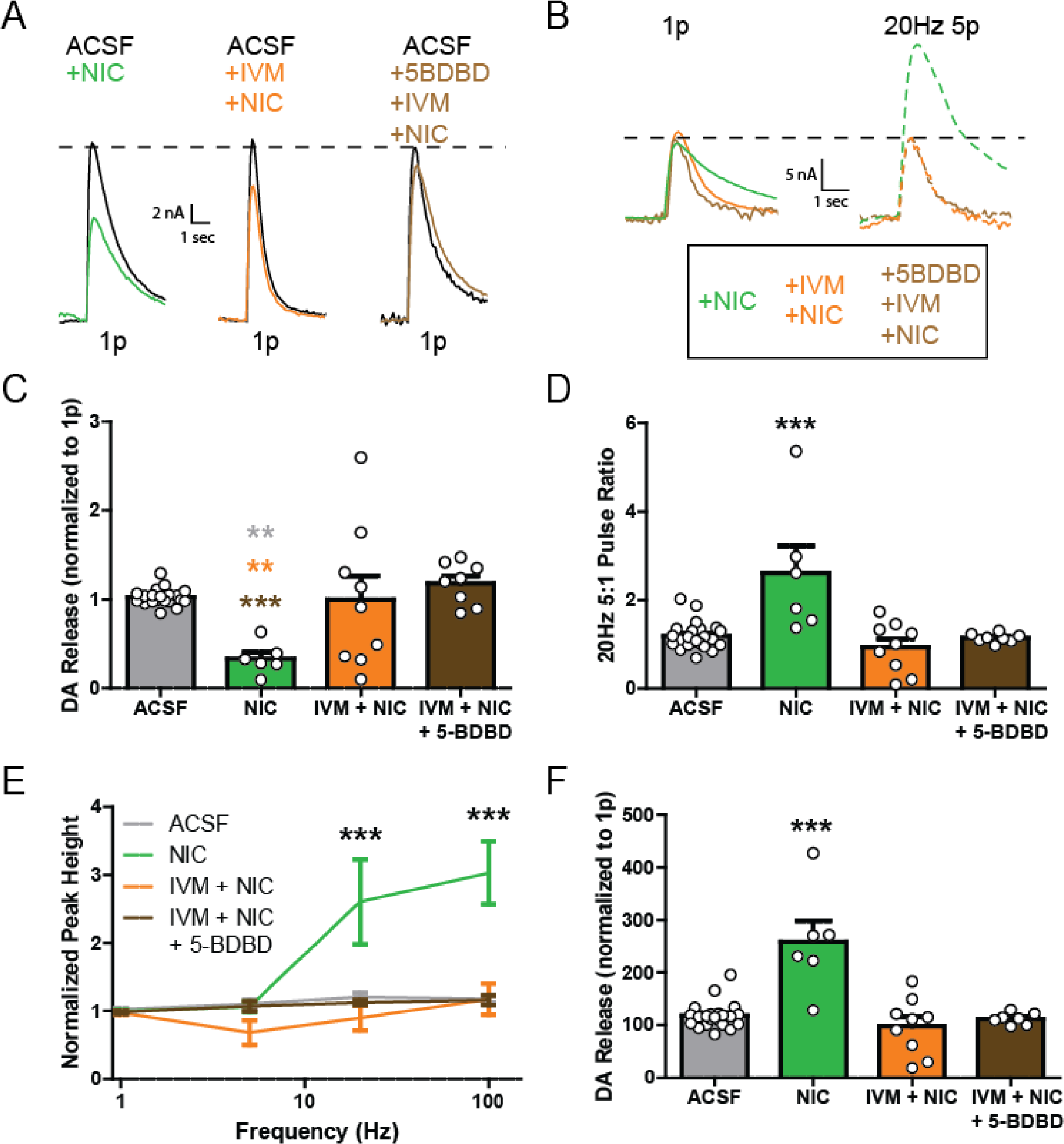
IVM attenuation of Nicotine effects is not through P2X4 receptors. **A** Representative traces of single pulse electrically evoked dopamine release in the DS. Nicotine (NIC) alone decreases single pulse dopamine release. **B** Representative traces of evoked dopamine release at multiple frequencies. Nicotine alone increases release with 20Hz 5 pulse stimulation compared to co-application with IVM or IVM and 5-BDBD. **C** Nicotine decreases single pulse dopamine release while IVM + NIC, and IVM + NIC + 5-BDBD have no effect on single pulse dopamine release compared to ACSF pre-treatment. **D** Nicotine alone increases the 20Hz 5 pulse ratio compared to ACSF, IVM + NIC, and IVM + NIC + 5-BDBD. **E** Nicotine alone shows increased dopamine release in response to high frequency, 5 pulse stimulations, with AUC represented in **F**. Asterisks ** indicate significance levels p < 0.01 compared to ACSF pre-treatment. Asterisks ** indicate significance levels p < 0.01 compared to IVM + NIC treatment. Asterisks *** indicate significance levels p < 0.001 compared to IVM + NIC + 5-BDBD. Asterisks *** indicate significance levels p < 0.001 compared to all other treatments.

### IVM enhances striatal cholinergic interneuron firing activity

Although IVM is known to interact with nicotinic receptors (Collins & Millar, 2010; Krause et al., 1998), IVM acetylcholine interactive effects may also involve indirect effects on CINs, and subsequent changes in acetylcholine release. Therefore, the effects of IVM on DS CIN firing was examined using cell-attached electrophysiology experiments. CINs were visually identified and patched onto using borosilicate glass capillary electrodes (**Fig. 4A**). IVM increased the CIN firing rate frequency from 1.32±0.53 to 3.15±1.47 Hz (**Fig. 4B**-**C**,**D**; Two-tailed Wilcoxon Matched Pairs test; *p* = 0.0156). The variance of firing frequency trended toward increases but was not significant (**Fig. 4E**; Two-tailed Wilcoxon Matched Pairs test; *p* = 0.1094). Thus, some of IVM effects are through excitatory effects on CIN firing, which are likely influencing the nicotine response observed in FSCV studies. Regardless, these data support the hypothesis that IVM is affecting DA release and nicotine effects through its effects on the striatal acetylcholine system.

**Figure 4:**
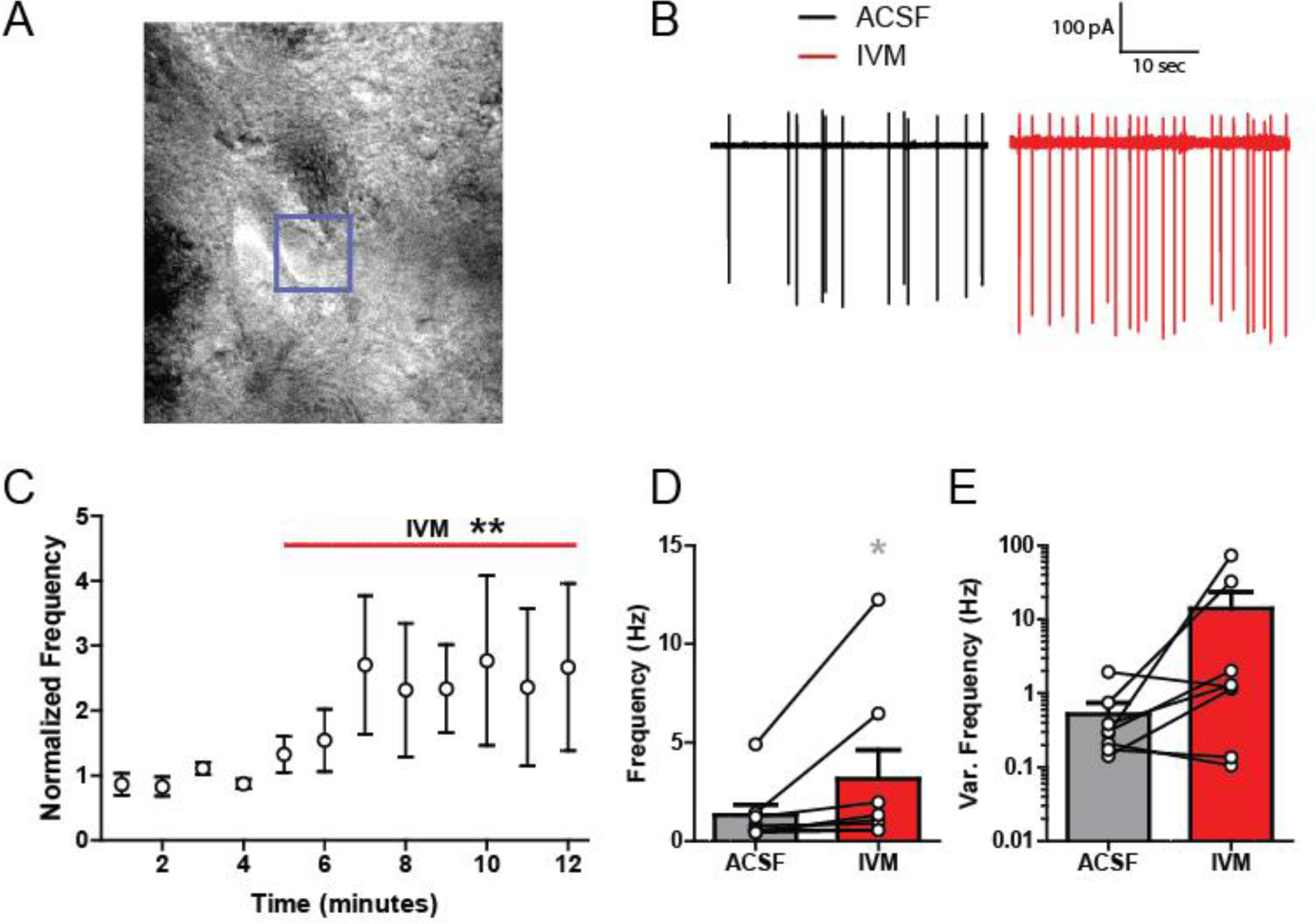
IVM effects on striatal cholinergic interneurons. **A** Representative image of glass capillary electrode connected to a DS CIN. **B** Representative traces of CIN firing rate before (left) and after IVM application (right). **C** Time course of CIN normalized frequency showing IVM is increasing DS CIN firing rate. **D** Ivermectin increases CIN firing rate frequency. **E** Action potential frequency variance (Var.) was increased in some but not all neurons, and group changes were not statistically significant. Asterisks * indicate significance levels p < 0.05 compared to ACSF pre-treatment.

### IVM effect on DA release is not through direct DA terminal activation

Whether IVM has direct effects on DA terminals independent of nAChRs was explored next. The nAChR antagonist hexamethonium (HEX; 200 µM) was applied to brain slices and interactions with IVM effects on DA release measured. Single pulse DA release was greatly reduced by HEX (**Fig. 5A**-**C**). When IVM was added to HEX, no noticeable change occurred for single pulse DA release, and normalized signals remained significantly lower compared to both baseline and IVM alone (**Fig. 5A**-**C**; one-way ANOVA; *F*_(3,60)_ = 84.14, p < 0.0001). Qualitatively, similar to that observed with nicotine, the 5:1 pulse ratio (20Hz) was increased after HEX, with no further increase observed with co-application of IVM + HEX (**Fig. 5D**; one-way ANOVA; *F*_(2,26)_ = 5.688, p = 0.0089). The frequency response for HEX and HEX+IVM were both significantly larger than pre-drug conditions, with normalized peak height becoming increasingly significant at higher frequencies (**Fig. 5E**; two-way ANOVA; drug, *F*_(2,117)_ = 14.85, p < 0.0001; frequency, *F*_(3,117)_ = 21.06, p < 0.0001; interaction, *F*_(3,117)_ = 3.229, p = 0.0057). In measuring the overall frequency response of each treatment using AUC, both HEX alone and HEX+IVM significantly increased the frequency effect compared to baseline levels (**Fig. 5F**; one-way ANOVA; *F*_(2,26)_ = 10.89, p = 0.0004). Together with agonist data from **Fig. 2**, the results in **Fig. 4** and **Fig. 5** support the hypothesis that IVM increases DA release through enhanced nicotinic receptor activity, and not through direct effects on DA terminals.

**Figure 5:**
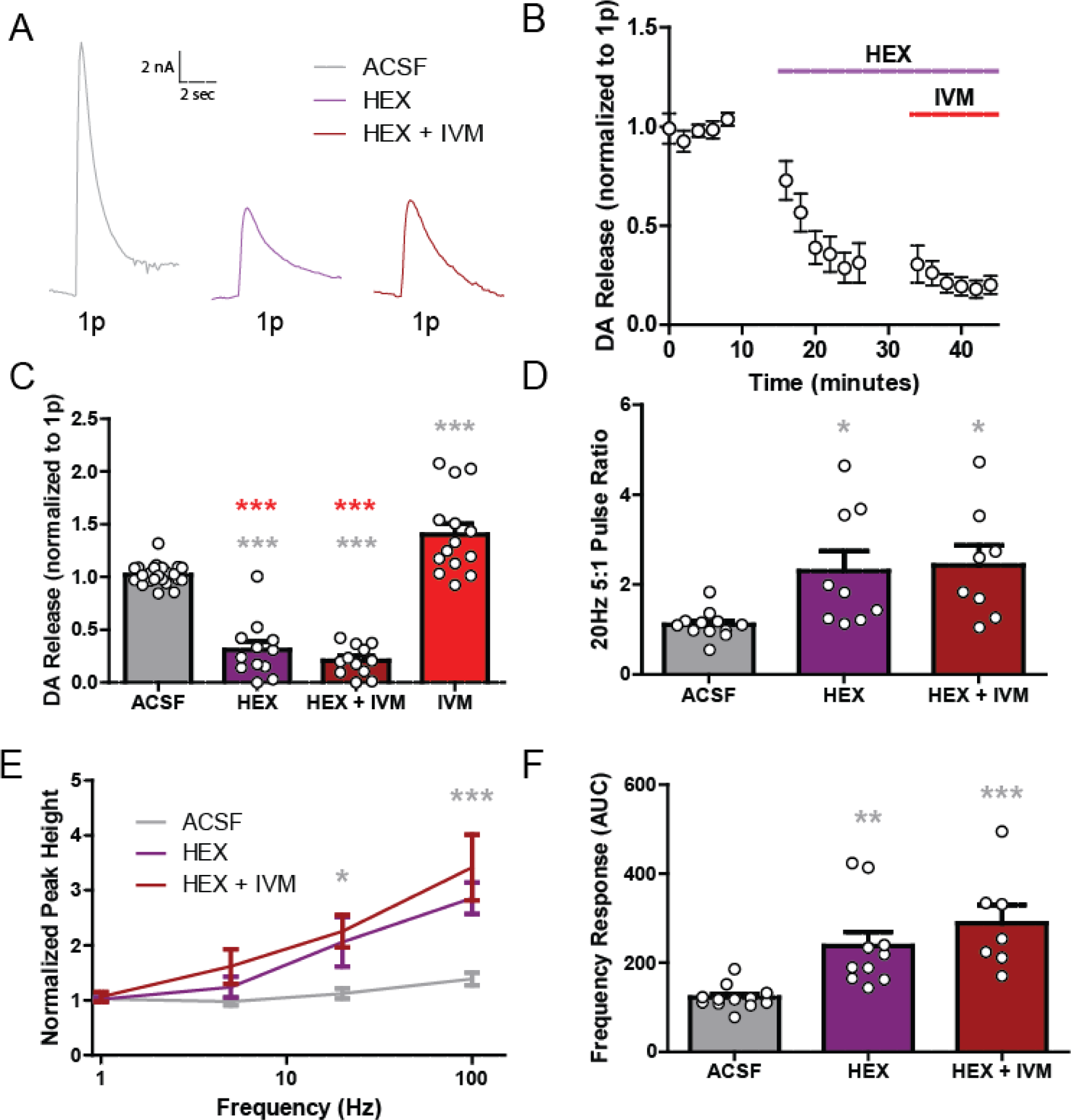
IVM effect on DA release is not through direct DA terminal activation. **A** Representative traces of single pulse electrical evoked dopamine release in the DS with HEX and HEX + IVM compared to ACSF pre-treatment. **B** Time course of single pulse dopamine release showing HEX induced decreases in dopamine release which is not rescued by concurrent IVM application. **C** Hexamethonium, and HEX + IVM decrease single pulse dopamine release compared to pre-treatment ACSF levels and to IVM, which increases single pulse dopamine release. **D** Hexamethonium and HEX + IVM increase the 20Hz 5 pulse ratio compared to ACSF pre-treatment. **E** Both HEX and HEX + IVM show increased dopamine release in response to high frequency, 5 pulse stimulations compared to ACSF pre-treatment. **F** Overall frequency response as measured by AUC. Asterisks *,**,*** indicate significance levels p < 0.05, p < 0.01, and p < 0.001, respectively compared to ACSF pre-treatment. Asterisks *** indicate significance levels p < 0.001 compared to IVM.

### IVM increases DA release with L-DOPA

Prior work has shown that IVM enhances L-DOPA induced behaviors(Warnecke et al., 2020). In consideration of this IVM and L-DOPA interaction, experiments were performed to test if IVM enhances L-DOPA effects on DA release. Application of L-DOPA increases single pulse DA release, which was significantly increased further with co-application of L-DOPA and IVM (**Fig. 6A**-**C**; one-way ANOVA; *F*_(3,83)_ = 22.74, p < 0.0001). Application of L-DOPA or IVM alone had no effect on the normalized 5:1 pulse ratio (20Hz) of DA release. However, combined application of L-DOPA with IVM statistically decreased the 5:1 pulse ratio compared to baseline levels (**Fig. 6D**; one-way ANOVA; *F*_(3,74)_ = 3.486, p = 0.0199). These results are reflective of paired pulse stimulation studies where enhanced release during the primary stimulation results in an attenuation in a secondary pulse(Manabe, Wyllie, Perkel, & Nicoll, 1993; Ronström et al., 2023). Therefore, the reduced 5:1 pulse ratio with IVM+L-DOPA co-application is likely due to increased release probability during the 1 pulse stimulation resulting in reduced readily releasable pools in the 5 pulse stimulation condition. This suggests that IVM increases DA release past what L-DOPA does alone, and that IVM effects are unique in mechanism to that of L-DOPA.

**Figure 6:**
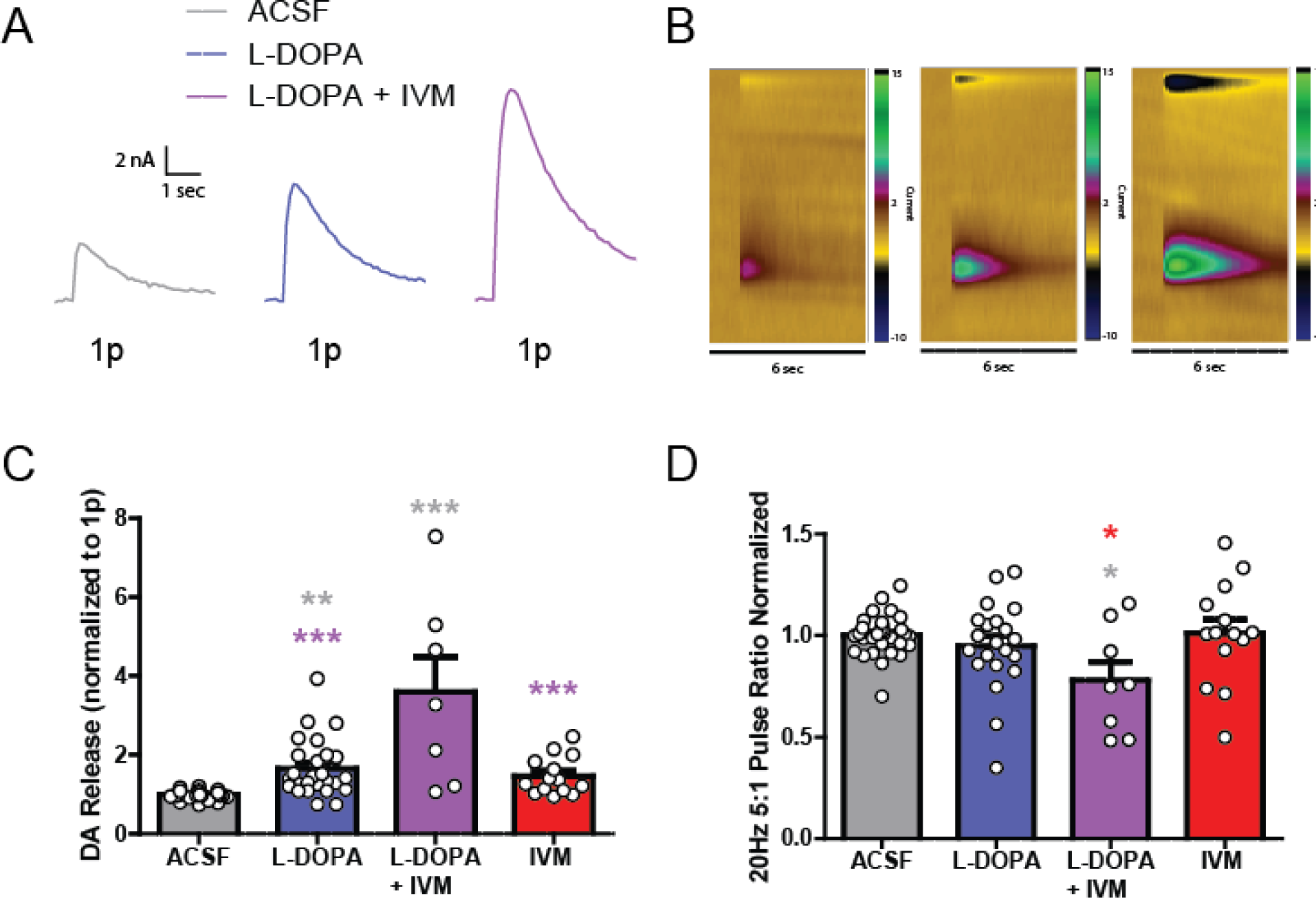
IVM increases DA release with L-DOPA. **A** Representative traces of single pulse electrical evoked dopamine release in the DS comparing pre-treatment ACSF (left), L-DOPA (middle) and L-DOPA + IVM (right). **B** Representative color plots of dopamine release with pre-treatment ACSF (left), L-DOPA (middle) and L-DOPA + IVM (right). **C** L-DOPA and L-DOPA + IVM significantly increases single pulse dopamine release compared to ACSF pre-treatment levels. **D** L-DOPA + IVM significantly decreases the normalized 20Hz 5 pulse ratio compared to ACSF pre-treatment, while L-DOPA alone and IVM alone have no effect. Asterisks *,**,*** indicate significance levels p < 0.05, p < 0.01, and p < 0.001, respectively compared to ACSF pre-treatment. Asterisks *** indicate significance levels p < 0.001 compared to L-DOPA + IVM. Asterisks * indicate significance levels p < 0.05 compared to IVM.

## Discussion

The present goal was to determine the effects of IVM on striatal DA release and pharmacological mechanisms. Ivermectin increased striatal DA release independent from P2X4 receptor activation. Furthermore, IVM reduced nicotine effects on DA release, and IVM effects were blocked by nicotinic receptor antagonism. Since IVM also enhanced CIN firing, IVM is likely influencing DA release through multiple cholinergic related mechanisms. Importantly, when IVM was co-applied with L-DOPA, there was a greater amount of DA release than with L-DOPA alone. L-DOPA is the biosynthetic precursor to DA and increases DA release by increasing the DA vesicular content(Omiatek et al., 2013). Ivermectin increases single pulse stimulation-mediated DA release but decreases the high frequency to low frequency DA release ratio when combined with L-DOPA. This decrease in ratio indicates that IVM effects are mostly through DA terminal excitation and not through additional changes in vesicular content.

Ivermectin has FDA approval and is used clinically to treat tropical parasitic diseases (Fox, 2006; Omura & Crump, 2014). While IVM was originally used for veterinary medicine in 1981, it was later approved for use in humans in 1988 (Crump & Ōmura, 2011), specifically for onchocerciasis, helminthiases and scabies (Omura & Crump, 2014). It is the only recommended oral medication for scabies (Gunning, Pippitt, Kiraly, & Sayler, 2012). Ivermectin works as an anti-parasitic by binding to invertebrate glutamate-gated chloride channels, causing hyperpolarization of parasite neurons and muscles leading to paralysis and eventually death (Gowtham & Karthikeyan, 2019). Ivermectin is also anti-inflammatory and relatively well tolerated (Kircik, Del Rosso, Layton, & Schauber, 2016). Because IVM is relatively safe, FDA approved, and commonly prescribed, it is important to understand its effects on neural circuitry. Furthermore, because it is an allosteric modulator, it has potential for amplifying pharmacological effects of other substances used clinically. Ivermectin itself increases DA release in the DS through cholinergic mechanisms, which could potentially benefit those with dysfunction in DA circuity, including those with PD, mood disorders, or attention deficit disorder.

### IVM effects through acetylcholine related mechanisms

Throughout the dorsal and ventral striatum, CINs are important frequency dependent modulators of DA release(Ferris, Calipari, Yorgason, & Jones, 2013; Nolan et al., 2020) and can drive DA release independent of DA cell firing(Wadsworth et al., 2023; Yorgason et al., 2017). Drugs that modulate CIN activity and nicotinic acetylcholine receptors on DA terminals enhance the ratio of DA release from high to low frequency stimulations, a well-characterized effect described as a high pass filter(Rice & Cragg, 2004; Sulzer, Cragg, & Rice, 2016; Zhang & Sulzer, 2004; Zhou et al., 2001). Despite the powerful regulation of DA by CINs, the present study did not indicate any potentiation of DA release under high frequency stimulations in the presence of IVM alone. This contrasted with IVM attenuating the nicotine-induced changes in DA release. This demonstrates that IVM is affecting the DS cholinergic system to modulate DA release. Since IVM directly increases CIN firing frequency, increases in DA release likely involve this change in CIN excitability and subsequent downstream effects on nAChRs. These findings were further verified by the application of HEX, which blocks CIN input onto DA terminals. By inhibiting the effects of acetylcholine through nAChR antagonism, IVM no longer increased single pulse DA release. Thus, IVM is likely not acting directly on DA terminals to enhance release. Interestingly, in the presence of IVM, HEX still induced a high-pass filter effect for multiple pulse stimulations. This suggests that nACHRs are the final common pathway for IVM effects, further strengthening the hypothesis that IVM is working through the cholinergic system and the communication between CIN and DA terminals.

### IVM effects on P2X4 receptors

Ivermectin is a PAM on P2X4 receptors, nicotinic receptors and γ-aminobutyric acid type A (GABA_A_) receptors(Bortolato et al., 2013; Coddou, Stojilkovic, & Huidobro-Toro, 2011; Crichlow, Mishra, & Crawford, 1986; Dawson et al., 2000; Diggs, Feller, Crabbe, Merrill, & Farrell, 1990; Jelínková et al., 2006; Khoja et al., 2016; Wyatt et al., 2014; Yardley et al., 2012). Notably, in P2X4 receptor knockout (KO) mice, the behavioral effects of IVM are diminished. Specifically, IVM decreases ethanol drinking and increases motor behavior, and these IVM effects are P2X4 receptor mediated(Khoja et al., 2016; Wyatt et al., 2014).

Presently, some of IVM’s effects on DA release were hypothesized to include the P2X4 receptor based on these prior studies. However, multiple experiments using P2X4 receptor antagonist 5-BDBD were performed, and P2X4 antagonism did not block any of IVM’s effects on DA release, including nicotine interactions. However, it’s possible that P2X4 receptors are involved in other aspects of these circuitry not tested herein. For instance, the P2X4 receptor is expressed on immune cells in the periphery, as well as microglia in the brain, which are activated through peripheral cytokine networks (Hoogland, Houbolt, van Westerloo, van Gool, & van de Beek, 2015). Furthermore, P2X4 receptor involvement could be present in other brain regions, including upstream circuits not present in brain slices. For instance, all present experiments were performed in DS brain slices, which is associated with motor behavior and decreases in DA release with PD(Shen, Zhai, & Surmeier, 2022). Therefore, the previous *in vivo* behavioral P2X4 KO studies have a peripheral immune component and additional circuitry components that were presently excluded. Previous work connecting P2X4 receptors to motor behaviors using P2X4 KO mice demonstrate that P2X4 receptors are involved in regulating DA terminal function(Khoja et al., 2016). Considering present experiments were only looking at acute IVM and 5-BDBD exposure, there may be effects of P2X4 receptors on DA release more directly over prolonged exposure. Regardless, current results provide strong evidence for IVM effects on DA terminals that are dependent on local cholinergic systems.

### L-DOPA

Previous studies in a mouse PD model showed that L-DOPA with IVM could alter DA motor behavior to a greater extent than L-DOPA on its own(Khoja et al., 2016; Warnecke et al., 2020). Presently, L-DOPA application increased DA release, which increased further with IVM co-application. The results of these experiments suggest that the behavioral effects seen in prior PD mouse models are due to L-DOPA and IVM’s ability to increase DA release through multiple mechanisms, specifically, increased vesicle DA content with L-DOPA(Qi et al., 2016), and increased release probability with IVM. Multimodal treatment approaches are common for PD(Van der Schyf & Geldenhuys, 2011). Ivermectin and L-DOPA interactions in humans have not been reported to date, though considering IVM and L-DOPA are both commonly prescribed, the lack of published studies suggests mild interactions at most for prescribed doses. Regardless, the present experiments suggest an interaction that may facilitate and/or complicate treatment. This was not further explored; L-DOPA was used as an experimental variable to test for mechanistic effects of IVM, and not to inform on PD treatment.

## Conclusion

This study provides novel information about the effects of IVM on DA release *in slice*. Dopamine (DA) release in the striatum is mediated by many intrinsic factors, including ion channels, autoreceptors and heteroreceptors. Through FSCV experiments, IVM is increasing DA release independent of P2X4 receptor activity in the DS, even though P2X4 receptors are in this region on microglia as well as other cells. Ivermectin also attenuates nicotine effects as a high-pass filter in a way that is not mediated through P2X4 receptors. In addition, IVM is able to increase CIN firing rate frequency, highlighting a possible mechanism through which IVM is acting to affect DA release. With that, HEX was used to block CIN outputs on nAChRs and saw that IVM no longer was able to increase DA release. Ivermectin was also unable to block HEX as a high pass filter, suggesting that IVM is not affecting DA release directly on DA terminals, providing further evidence of IVM acting on DA release through CINs. And when IVM was co-applied with L-DOPA there was an even greater increase of DA release compared to IVM or L-DOPA alone. In addition, co-application of L-DOPA with IVM saw a decrease in the normalized 5 pulse ratio, showing that L-DOPA and IVM together is decreasing the readily releasable pool of DA, meaning that IVM is acting through a different mechanism then L-DOPA and highlighting IVM is not acting on DA terminals. This study has helped elucidate the effects of IVM on DA release and how it is able to mediate the cholinergic system in the DS.

## Declaration of Interest

There are no declarations of interests to declare.

## Notes

### Competing Interest Statement

The authors have declared no competing interest.

